# The impact of the ABO/Rh blood group on susceptibility and severity among COVID-19 patients in Luanda, Angola

**DOI:** 10.1101/2022.10.14.512325

**Authors:** Cruz S. Sebastião, Alice Teixeira, Ana Luísa, Margarete Arrais, Chissengo Tchonhi, Adis Cogle, Euclides Sacomboio, Bruno Cardoso, Joana Morais, Jocelyne Neto de Vasconcelos, Miguel Brito

## Abstract

SARS-CoV-2 is a public health concern worldwide. Identification of biological factors that could influence transmission and worsen the disease has been the subject of extensive investigation. Herein, we investigate the impact of the ABO/Rh blood group on susceptibility and severity among COVID-19 patients in Luanda, Angola. This was a multicentric cohort study conducted with 101 COVID-19 patients. Chi-square and logistic regression were calculated to check factors related to the worsening of the disease and deemed significant when p<0.05. Blood type O (51.5%) and Rh-positive (93.1%) were the most frequent. Patients from blood type O had a high risk to severe disease [OR: 1.33 (95% CI: 0.42 - 4.18), p=0.630] and hospitalization [OR: 2.59 (95% CI: 0.84 - 8.00), p=0.099]. Also, Rh-positive blood type presented a high risk for severe disease (OR: 10.6, p=0.007) and hospitalization (OR: 6.04, p=0.026). We find a high susceptibility, severity, hospitalization, and mortality, respectively, among blood group O and Rh-positive patients, while blood group AB presented a low susceptibility, severity, hospitalization, and mortality, respectively. Our findings add to the body of evidence suggesting that ABO/Rh blood groups play an important role in the course of SARS-CoV-2 infection.

## Introduction

Severe acute respiratory syndrome coronavirus 2 (SARS-CoV-2) was initially identified in Wuhan, one of China’s provinces at the end of 2019, quickly spread and evolved into a global public health emergency. By the end of June 2021, more than 180 million cases and 3.9 million deaths were related to SARS-CoV-2. During the same period, Angola recorded more than 39,000 cases and 920 deaths related to Coronavirus disease (COVID-19). [1] COVID-19-related mortality rates have been driven by patients who develop respiratory failure during the SARS-CoV-2 infection. [2] Therefore, the pathogenesis of respiratory failure in COVID-19 patients is unclear, although studies indicate that mortality is associated with older age, male gender, and comorbidities such as hypertension, diabetes, cardiovascular disease, and obesity. [3]

Susceptibility of certain viral infections and diseases has been linked to ABO/Rh blood group polymorphism. [4–6] Previous studies have shown that ABO blood group polymorphisms increase the COVID-19 severity among blood group A patients and reduce the severity in blood group O patients. [7–10] Thereby, the different ABO human blood groups have been used as important biomarkers for disease prediction. [11] Currently, studying the relationship between ABO blood groups and SARS-CoV-2 infection can be crucial for determining the vulnerability of infection in the population and help define strategies for immediate clinical intervention, especially in low- and middle-income countries (LMICs) where the COVID-19 pandemic response may be threatened due to limited resources. [12,13] There is a growing interest in identifying the possible risk factors that determine vulnerability to infection or worsening of the clinical condition among COVID-19 patients. Currently, there is no published study showing the impact of blood group polymorphisms as well as their relationship to SARS-CoV-2 infection and severity in Angola. In this study, we investigate for the first time, the impact of the ABO/Rh blood group on susceptibility and severity among COVID-19 patients in Luanda, the capital city of Angola, to contribute to global knowledge about SARS-CoV-2 infection and to support the management of the COVID-19 patients in Angola.

## Materials and methods

### Study design and setting

A multicentric cohort study was carried out with 101 subjects infected with SARS-CoV-2 at the Hospital Militar Principal, Clínica Girassol, and at the Lucrécia Paim maternity, all located in Luanda, between December 2020 to April 2021. The study was approved by the national ethics committee of the Ministry of Health of Angola (approval nr. 25/2020), the general director of Hospital Militar Principal (approval nr. 2193/DDI/HMP/IS/20), the general director of Clínica Girassol (approval nr. 1945/GEPP/PCE/2020), and general director of Lucrécia Paim maternity (approval nr. 840/GDG/MLP/2020). Participants were informed of the study and verbal consent was obtained from participants before being included in the study.

### Sample collection and testing

A structured questionnaire was used to collect sociodemographic (age, gender, and residence area) and clinical (symptoms, comorbidities, and clinic category) data. Only individuals with positive SARS-CoV-2 infection were included in the study, whereas individuals with negative or inconclusive results for SARS-CoV-2 infection were excluded from the study. The SARS-CoV-2 infection was screened and confirmed by at least one quantitative real-time reverse transcriptase-polymerase chain reaction (qRT-PCR) assay with the Applied Biosystems 7500 Fast RT-PCR System (Thermo Fisher Scientific), using a protocol previously described for the detection of 2019 novel Coronavirus (2019-nCoV) RNA (PCR-Fluorescence Probing) (Da An Gene, China). [14] Patients were followed up on clinical outcome and all with a negative RT-PCR result during the follow-up period were considered to recover with loss of follow-up. An estimated volume of 3 mL of whole blood was collected in a tube containing EDTA for the determination of ABO/RH blood group phenotypes (Lorne Laboratories Limited, UK), following the manufacturer’s instructions. [15] Lorne Monoclonal IgM ABO blood grouping reagents contain mouse monoclonal antibodies diluted in a phosphate buffer containing sodium chloride, EDTA, and bovine albumin. Each reagent is supplied at optimal dilution for use by slide, tube, gel card, and microplate techniques. [15] The laboratory procedures for the determination of ABO/Rh blood groups were performed in the hemotherapy of Clínica Girassol and the hematology laboratory of INIS, both located in Luanda.

### Statistical analysis

The statistical analysis was carried out using SPSS version 26 (IBM SPSS Statistics, USA). Frequencies and percentages were presented as descriptive analyses. Normally distributed data were presented as mean and standard deviation. Chi-square (X^2^) test and univariate logistic regression analysis were performed to check interactions between categorical variables. Odds ratio (OR) with their 95% confidence intervals (CIs) were also calculated to determine the strength of the interaction between variables and were deemed significant when p<0.05.

## Results

### Sociodemographic and clinical characteristics related to ABO/Rh blood groups

The sociodemographic and clinical characteristics related to ABO/Rh blood groups among COVID-19 patients in Luanda are summarized in Table 1. This study included a total of 101 patients diagnosed with SARS-CoV-2 infection by RT-PCR in Luanda, between December 2020 – April 2021. Overall, blood group O (51.5%, 52/101) was the most frequent, followed by blood groups A (24.8%, 25/101), B (20.8%, 21/101), and AB (3%, 3/101). The positive RH factor predominated with 93.1% (94/101) compared to the negative RH factor (6.9%, 7/101). Age ranged from 18 to 80 years. The mean age was 51±14 years old. Patients aged over 40 years (75.2%, 76/101), male (60.4%, 61/101), and living in urbanized areas (54.5%, 55/101), were predominant in this study population. Clinical characteristics showed that 77.2% (78/101) of patients had symptoms related to COVID-19, with 54.5% (55/101) having a moderate SARS-CoV-2 infection, followed by patients with mild and/or severe infections with 22.8% (23/101), simultaneously. In addition, 64.4% (65/101) of COVID-19 patients had different comorbidities (such as hypertension, diabetes, kidney disease, obesity, and cerebrovascular accident), 79.2% (80/101) required hospitalization, and 9.9% (10/101) died due to complications related to COVID-19. The hospitalization rate was between 2.5% (group AB) to 55% (group O), however, only patients in group A (8%, 2/25) and group O (15.4, 8/52) died as a clinical outcome. A significant relationship was observed among blood group A with the presence of fever (p=0.049), while blood group AB was related to malaise (p=0.039), anorexia (0.042), and malaria (p=0.002). On the other hand, the Rh blood type was significantly related to clinical category (p=0.005), hospitalization (p=0.014), and clinical symptoms (p=0.001).

**Table 1.** Sociodemographic and clinical characteristics related to ABO/Rh blood types among COVID-19 patients in Luanda, Angola.

| Characteristics | N (%) | ABO blood group distribution |  |  |  |  |  |  |  |  |  |  | Rh blood type |  |  |  |
| --- | --- | --- | --- | --- | --- | --- | --- | --- | --- | --- | --- | --- | --- | --- | --- | --- |
|  |  | A |  |  | B |  |  | AB |  |  | O |  |  |  |  |  |
|  |  | No (%) | Yes (%) | p-value | No (%) | Yes (%) | p-value | No (%) | Yes (%) | p-value | No (%) | Yes (%) | p-value | Neg. (%) | Pos. (%) | p-value |
| Overall | 101 (100) | 76 (75.2) | 25 (24.8) |  | 80 (79.2) | 21 (20.8) |  | 98 (97.0) | 3 (3.0) |  | 49 (48.5) | 52 (51.5) |  | 7 (6.90) | 94 (93.1) |  |
| Age groups |  |  |  |  |  |  |  |  |  |  |  |  |  |  |  |  |
| <20y | 1 (1.0) | 0 (0.0) | 1 (100) | 0.070 | 1 (100) | 0 (0.0) | 0.732 | 1 (100) | 0 (0.0) | 0.601 | 1 (100) | 0 (0.0) | 0.062 | 0 (0.0) | 1 (100) | 0.791 |
| 20 – 40y | 24 (23.8) | 21 (87.5) | 3 (12.5) |  | 20 (83.3) | 4 (16.7) |  | 24 (100) | 0 (0.0) |  | 7 (29.2) | 17 (70.8) |  | 1 (4.20) | 23 (95.8) |  |
| >40y | 76 (75.2) | 55 (72.4) | 21 (27.6) |  | 59 (77.6) | 17 (22.4) |  | 73 (96.1) | 3 (3.90) |  | 41 (53.9) | 35 (46.1) |  | 6 (7.90) | 70 (92.1) |  |
| Gender |  |  |  |  |  |  |  |  |  |  |  |  |  |  |  |  |
| Female | 40 (39.6) | 29 (72.5) | 11 (27.5) | 0.604 | 35 (87.5) | 5 (12.5) | 0.096 | 40 (100) | 0 (0.0) | 0.154 | 16 (40.0) | 24 (60.0) | 0.166 | 2 (5.0) | 38 (95.0) | 0.536 |
| Male | 61 (60.4) | 47 (77.0) | 14 (23.0) |  | 45 (73.8) | 16 (26.2) |  | 58 (95.1) | 3 (4.90) |  | 33 (54.1) | 28 (45.9) |  | 5 (8.20) | 56 (91.8) |  |
| Residence area |  |  |  |  |  |  |  |  |  |  |  |  |  |  |  |  |
| Rural | 46 (45.5) | 36 (78.3) | 10 (21.7) | 0.521 | 34 (73.9) | 12 (26.1) | 0.230 | 46 (100) | 0 (0.0) | 0.108 | 22 (47.8) | 24 (52.2) | 0.899 | 2 (4.30) | 44 (95.7) | 0.350 |
| Urban | 55 (54.5) | 40 (72.7) | 15 (27.3) |  | 46 (83.6) | 9 (16.4) |  | 52 (94.5) | 3 (5.50) |  | 27 (49.1) | 28 (50.9) |  | 5 (9.10) | 50 (90.9) |  |
| Clinical category |  |  |  |  |  |  |  |  |  |  |  |  |  |  |  |  |
| Mild | 23 (22.8) | 17 (73.9) | 6 (26.1) | 0.714 | 17 (73.9) | 6 (26.1) | 0.531 | 22 (95.7) | 1 (4.30) | 0.625 | 13 (56.5) | 10 (43.5) | 0.651 | 5 (21.7) | 18 (78.3) | 0.005 |
| Moderate | 55 (54.5) | 43 (78.2) | 12 (21.8) |  | 43 (78.2) | 12 (21.8) |  | 53 (96.4) | 2 (3.60) |  | 26 (47.3) | 29 (52.7) |  | 2 (3.60) | 53 (96.4) |  |
| Severe | 23 (22.8) | 16 (69.6) | 7 (30.4) |  | 20 (87.0) | 3 (13.0) |  | 23 (100) | 0 (0.0) |  | 10 (43.5) | 13 (56.5) |  | 0 (0.0) | 23 (100) |  |
| Hospitalization |  |  |  |  |  |  |  |  |  |  |  |  |  |  |  |  |
| No | 21 (20.8) | 13 (61.9) | 8 (38.1) | 0.111 | 17 (81.0) | 4 (19.0) | 0.825 | 20 (95.2) | 1 (4.80) | 0.587 | 13 (61.9) | 8 (38.1) | 0.168 | 4 (19.0) | 17 (81.0) | 0.014 |
| Yes | 80 (79.2) | 63 (78.8) | 17 (21.3) |  | 63 (78.8) | 17 (21.3) |  | 78 (97.5) | 2 (2.50) |  | 36 (45.0) | 44 (55.0) |  | 3 (3.80) | 77 (96.3) |  |
| Clinical outcome |  |  |  |  |  |  |  |  |  |  |  |  |  |  |  |  |
| Recovery | 91 (90.1) | 68 (74.7) | 23 (25.3) | 0.714 | 70 (76.9) | 21 (23.1) | 0.088 | 88 (96.7) | 3 (3.30) | 0.560 | 47 (51.6) | 44 (48.4) | 0.057 | 7 (7.70) | 84 (92.3) | 0.363 |
| Death | 10 (9.90) | 8 (80.0) | 2 (20.0) |  | 10 (100) | 0 (0.0) |  | 10 (100) | 0 (0.0) |  | 2 (20.0) | 8 (80.0) |  | 0 (0.0) | 10 (100) |  |
| Symptoms |  |  |  |  |  |  |  |  |  |  |  |  |  |  |  |  |
| No | 23 (22.8) | 17 (73.9) | 6 (26.1) | 0.866 | 17 (73.9) | 6 (26.1) | 0.476 | 22 (95.7) | 1 (4.30) | 0.658 | 13 (56.5) | 10 (43.5) | 0.382 | 5 (21.7) | 18 (78.3) | 0.001 |
| Yes | 78 (77.2) | 59 (75.6) | 19 (24.4) |  | 63 (80.8) | 15 (19.2) |  | 76 (97.4) | 2 (2.60) |  | 36 (46.2) | 42 (53.8) |  | 2 (2.60) | 76 (97.4) |  |
| Fever |  |  |  |  |  |  |  |  |  |  |  |  |  |  |  |  |
| No | 65 (64.4) | 53 (81.5) | 12 (18.5) | 0.049 | 49 (75.4) | 16 (24.6) | 0.203 | 64 (98.5) | 1 (1.50) | 0.255 | 29 (44.6) | 36 (55.4) | 0.292 | 6 (9.20) | 59 (90.8) | 0.221 |
| Yes | 36 (35.6) | 23 (63.9) | 13 (36.1) |  | 31 (86.1) | 5 (13.9) |  | 34 (94.4) | 2 (5.60) |  | 20 (55.6) | 16 (44.4) |  | 1 (2.80) | 35 (97.2) |  |
| Cough |  |  |  |  |  |  |  |  |  |  |  |  |  |  |  |  |
| No | 64 (63.4) | 50 (78.1) | 14 (21.9) | 0.378 | 49 (76.6) | 15 (23.4) | 0.389 | 61 (95.3) | 3 (4.70) | 0.181 | 32 (50.0) | 32 (50.0) | 0.694 | 5 (7.80) | 59 (92.2) | 0.646 |
| Yes | 37 (36.6) | 26 (70.3) | 11 (29.7) |  | 31 (83.8) | 6 (16.2) |  | 37 (100) | 0 (0.0) |  | 17 (45.9) | 20 (54.1) |  | 2 (5.40) | 35 (94.6) |  |
| Fatigue |  |  |  |  |  |  |  |  |  |  |  |  |  |  |  |  |
| No | 93 (92.1) | 69 (74.2) | 24 (25.8) | 0.403 | 73 (78.5) | 20 (21.5) | 0.547 | 91 (97.8) | 2 (2.20) | 0.098 | 46 (49.5) | 47 (50.5) | 0.516 | 7 (7.50) | 86 (92.5) | 0.421 |
| Yes | 8 (7.90) | 7 (87.5) | 1 (12.5) |  | 7 (87.5) | 1 (12.5) |  | 7 (87.5) | 1 (12.5) |  | 3 (37.5) | 5 (62.5) |  | 0 (0.0) | 8 (100) |  |
| Dyspnoea |  |  |  |  |  |  |  |  |  |  |  |  |  |  |  |  |
| No | 82 (81.2) | 61 (74.4) | 21 (25.6) | 0.678 | 65 (79.3) | 17 (20.7) | 0.975 | 80 (97.6) | 2 (2.40) | 0.514 | 40 (48.8) | 42 (51.2) | 0.912 | 6 (7.30) | 76 (92.7) | 0.751 |
| Yes | 19 (18.8) | 15 (78.9) | 4 (21.1) |  | 15 (78.9) | 4 (21.1) |  | 18 (94.7) | 1 (5.30) |  | 9 (47.4) | 10 (52.6) |  | 1 (5.30) | 18 (94.7) |  |
| Osteomyotralgia |  |  |  |  |  |  |  |  |  |  |  |  |  |  |  |  |
| No | 85 (84.2) | 62 (72.9) | 23 (27.1) | 0.216 | 67 (78.8) | 18 (21.2) | 0.826 | 83 (97.6) | 2 (2.40) | 0.400 | 43 (50.6) | 42 (49.4) | 0.337 | 7 (8.20) | 78 (91.8) | 0.234 |
| Yes | 16 (15.8) | 14 (87.5) | 2 (12.5) |  | 13 (81.3) | 3 (18.8) |  | 15 (93.8) | 1 (6.30) |  | 6 (37.5) | 10 (62.5) |  | 0 (0.0) | 16 (100) |  |
| Headache |  |  |  |  |  |  |  |  |  |  |  |  |  |  |  |  |
| No | 86 (85.1) | 63 (73.3) | 23 (26.7) | 0.267 | 69 (80.2) | 17 (19.8) | 0.543 | 83 (96.5) | 3 (3.50) | 0.463 | 43 (50.0) | 43 (50.0) | 0.475 | 7 (8.10) | 79 (91.9) | 0.252 |
| Yes | 15 (14.9) | 13 (86.7) | 2 (13.3) |  | 11 (73.3) | 4 (26.7) |  | 15 (100) | 0 (0.0) |  | 6 (40.0) | 9 (60.0) |  | 0 (0.0) | 15 (100) |  |
| Vomit |  |  |  |  |  |  |  |  |  |  |  |  |  |  |  |  |
| No | 97 (96.0) | 74 (76.3) | 23 (23.7) | 0.233 | 76 (78.4) | 21 (21.6) | 0.296 | 94 (96.9) | 3 (3.10) | 0.721 | 47 (48.5) | 50 (51.5) | 0.952 | 7 (7.20) | 90 (92.8) | 0.578 |
| Yes | 4 (4.00) | 2 (50.0) | 2 (50.0) |  | 4 (100) | 0 (0.0) |  | 4 (100) | 0 (0.0) |  | 2 (50.0) | 2 (50.0) |  | 0 (0.0) | 4 (100) |  |
| Diarrhea |  |  |  |  |  |  |  |  |  |  |  |  |  |  |  |  |
| No | 96 (95.0) | 74 (77.1) | 22 (22.9) | 0.061 | 75 (78.1) | 21 (21.9) | 0.240 | 93 (96.9) | 3 (3.10) | 0.688 | 46 (47.9) | 50 (52.1) | 0.598 | 6 (6.30) | 90 (93.8) | 0.238 |
| Yes | 5 (5.00) | 2 (40.0) | 3 (60.0) |  | 5 (100) | 0 (0.0) |  | 5 (100) | 0 (0.0) |  | 3 (60.0) | 2 (40.0) |  | 1 (20.0) | 4 (80.0) |  |
| Malaise |  |  |  |  |  |  |  |  |  |  |  |  |  |  |  |  |
| No | 81 (80.2) | 61 (75.3) | 20 (24.7) | 0.977 | 61 (75.3) | 20 (24.7) | 0.052 | 80 (98.8) | 1 (1.20) | **0.039** | 41 (50.6) | 40 (49.4) | 0.395 | 6 (7.40) | 75 (92.6) | 0.704 |
| Yes | 20 (19.8) | 15 (75.0) | 5 (25.0) |  | 19 (95.0) | 1 (5.0) |  | 18 (90.0) | 2 (10.0) |  | 8 (40.0) | 12 (60.0) |  | 1 (5.0) | 19 (95.0) |  |
| Asthenia |  |  |  |  |  |  |  |  |  |  |  |  |  |  |  |  |
| No | 74 (73.3) | 55 (74.3) | 19 (25.7) | 0.722 | 59 (79.7) | 15 (20.3) | 0.831 | 73 (98.6) | 1 (1.40) | 0.113 | 35 (47.3) | 39 (52.7) | 0.685 | 5 (6.80) | 69 (93.2) | 0.909 |
| Yes | 27 (26.7) | 21 (77.8) | 6 (22.2) |  | 21 (77.8) | 6 (22.2) |  | 25 (92.6) | 2 (7.40) |  | 14 (51.9) | 13 (48.1) |  | 2 (7.40) | 25 (92.6) |  |
| Anorexia |  |  |  |  |  |  |  |  |  |  |  |  |  |  |  |  |
| No | 95 (94.1) | 71 (74.7) | 24 (25.3) | 0.636 | 77 (81.1) | 18 (18.9) | 0.069 | 93 (97.9) | 2 (2.10) | **0.042** | 44 (46.3) | 51 (53.7) | 0.078 | 7 (7.40) | 88 (92.6) | 0.491 |
| Yes | 6 (5.90) | 5 (83.3) | 1 (16.7) |  | 3 (50.0) | 3 (50.0) |  | 5 (83.3) | 1 (16.7) |  | 5 (83.3) | 1 (16.7) |  | 0 (0.0) | 6 (100) |  |
| Anosmia |  |  |  |  |  |  |  |  |  |  |  |  |  |  |  |  |
| No | 92 (91.1) | 69 (75.0) | 23 (25.0) | 0.854 | 72 (78.3) | 20 (21.7) | 0.453 | 90 (97.8) | 2 (2.20) | 0.132 | 45 (48.9) | 47 (51.1) | 0.798 | 7 (7.60) | 85 (92.4) | 0.391 |
| Yes | 9 (8.90) | 7 (77.8) | 2 (22.2) |  | 8 (88.9) | 1 (11.1) |  | 8 (88.9) | 1 (11.1) |  | 4 (44.4) | 5 (55.6) |  | 0 (0.0) | 9 (100) |  |
| Hemiplegia |  |  |  |  |  |  |  |  |  |  |  |  |  |  |  |  |
| No | 100 (99.0) | 75 (75.0) | 25 (25.0) | 0.564 | 79 (79.0) | 21 (21.0) | 0.607 | 97 (97.0) | 3 (3.0) | 0.860 | 49 (49.0) | 51 (51.0) | 0.329 | 7 (7.0) | 93 (93.0) | 0.784 |
| Yes | 1 (1.00) | 1 (100) | 0 (0.0) |  | 1 (100) | 0 (0.0) |  | 1 (100) | 0 (0.0) |  | 0 (0.0) | 1 (100) |  | 0 (0.0) | 1 (100) |  |
| Consciousness loss |  |  |  |  |  |  |  |  |  |  |  |  |  |  |  |  |
| No | 100 (99.0) | 75 (75.0) | 25 (25.0) | 0.564 | 79 (79.0) | 21 (21.0) | 0.607 | 97 (97.0) | 3 (3.0) | 0.860 | 49 (49.0) | 51 (51.0) | 0.329 | 7 (7.0) | 93 (93.0) | 0.784 |
| Yes | 1 (1.00) | 1 (100) | 0 (0.0) |  | 1 (100) | 0 (0.0) |  | 1 (100) | 0 (0.0) |  | 0 (0.0) | 1 (100) |  | 0 (0.0) | 1 (100) |  |
| Comorbidities<sup>#</sup> |  |  |  |  |  |  |  |  |  |  |  |  |  |  |  |  |
| No | 36 (35.6) | 26 (72.2) | 10 (27.8) | 0.600 | 26 (72.2) | 10 (27.8) | 0.198 | 35 (97.2) | 1 (2.80) | 0.932 | 21 (58.3) | 15 (41.7) | 0.142 | 3 (8.30) | 33 (91.7) | 0.680 |
| Yes | 65 (64.4) | 50 (76.9) | 15 (23.1) |  | 54 (83.1) | 11 (16.9) |  | 63 (96.9) | 2 (3.10) |  | 28 (43.1) | 37 (56.9) |  | 4 (6.20) | 61 (93.8) |  |
| Malaria |  |  |  |  |  |  |  |  |  |  |  |  |  |  |  |  |
| No | 98 (97.0) | 75 (76.5) | 23 (23.5) | 0.088 | 77 (78.6) | 21 (21.4) | 0.368 | 96 (98.0) | 2 (2.0) | **0.002** | 46 (46.9) | 52 (53.1) | 0.070 | 7 (7.10) | 91 (92.9) | 0.631 |
| Yes | 3 (3.00) | 1 (33.3) | 2 (66.7) |  | 3 (100) | 0 (0.0) |  | 2 (66.7) | 1 (33.3) |  | 3 (100) | 0 (0.0) |  | 0 (0.0) | 3 (100) |  |

Abbreviations: Neg, Negative; Pos, Positive
Bold numbers mean that results were statistically significant for the X² test (p<0.05).
<sup>#</sup>Comorbidities: Hypertension; Diabetes; Kidney disease; Obesity; Cerebrovascular accident.

### Relationship between ABO/Rh blood groups with disease severity and hospitalization

The putative relationship between ABO/Rh blood groups with disease severity and hospitalization among COVID-19 patients from Luanda is summarized in Table 2. Just a few statistically significant associations were observed. Patients of blood group O had a high chance of developing symptomatic SARS-CoV-2 infection [OR: 1.33 (95% CI: 0.42 - 4.18), p=0.630] and hospitalization [OR: 2.59 (95% CI: 0.84 - 8.00), p=0.099]. A reduced chance of developing symptomatic infection [OR: 0.66 (95% CI: 0.26 - 1.68), p=0.384] as well as hospitalization [OR: 0.50 (95% CI: 0.19 - 1.35), p=0.172] was observed among non-O blood group patients, when compared to patients from blood group O. Blood group AB patients had a reduced chance of having symptomatic SARS-CoV-2 infection [OR: 0.63 (95% CI: 0.05 – 8.25), p=0.632] and hospitalization [OR: 0.94 (95% CI: 0.07 – 12.0), p=0.963]. Patients from non-AB blood groups had high chances of developing symptomatic infection [OR: 1.73 (95% CI: 0.15 - 20.0), p=0.662] and hospitalization [OR: 1.95 (95% CI: 0.17 - 22.6), p=0.593], when compared to patients from blood group AB. Blood group B patients had a reduced chance of having symptomatic SARS-CoV-2 infection [OR: 0.79 (95% CI: 0.21 – 2.95), p=0.725], although the same group had a high risk of hospitalization [OR: 2.0 (95% CI: 0.51 – 7.92), p=0.323]. Non-B blood group patients have a high chance of developing symptomatic infection [OR: 1.48 (95% CI: 0.50 - 4.40), p=0.478], but the same group has a reduced risk of hospitalization [OR: 0.87 (95% CI: 0.26 - 2.94), p=0.825] when compared to patients from blood group B. The non-A blood groups are more likely to develop symptomatic infection [OR: 1.10 (95% CI: 0.38 – 3.18), p=0.866] and to be hospitalized for worsening infection [OR: 2.28 (95% CI: 0.81 - 6.39), p=0.117], compared to blood group A. Regarding Rh blood type, our results showed high chances of developing symptomatic SARS-CoV-2 infection [OR: 10.6 (95% CI: 1.89 - 58.9), p=0.007] and hospitalization [OR: 6.04 (95% CI: 1.24 - 29.5), p=0.026] among Rh-positive patients when compared to Rh- negative patients. Interestingly, a low risk of developing symptomatic infection and hospitalization was observed in all Rh-negative blood groups among patients of the same blood group (eg, A+ vs. A-, B+ vs. B-, and O+ vs. O−) or in patients from different blood groups (eg, A+ vs. B-, A+ vs. O−, B+ vs. O−, AB+ vs. B-, AB+ vs. O−, and O+ vs. B-), although we did not observe statistical significance (p>0.05).

**Table 2.** Relationship between ABO/Rh blood groups with disease severity and hospitalization among COVID-19 patients in Luanda, Angola.

| Blood group distribution | All patients (N=101) | Symptomatic disease |  | Hospitalization |  |
| --- | --- | --- | --- | --- | --- |
|  |  | OR (95% CI) | p-value | OR (95% CI) | P-value |
| ABO phenotype |  |  |  |  |  |
| A | 25 (24.8) | 1.00 | - | 1.00 | - |
| B | 21 (20.8) | 0.79 (0.21 – 2.95) | 0.725 | 2.00 (0.51 – 7.92) | 0.323 |
| AB | 3 (3.00) | 0.63 (0.05 – 8.25) | 0.632 | 0.94 (0.07 – 12.0) | 0.963 |
| O | 52 (51.5) | 1.33 (0.42 – 4.18) | 0.630 | 2.59 (0.84 – 8.00) | 0.099 |
| ABO phenotype comparison |  |  |  |  |  |
| A vs. B | 25 (24.8); 21 (20.8)] | 0.79 (0.21 – 2.95) | 0.725 | 2.00 (0.51 – 7.92) | 0.323 |
| A vs. AB | 25 (24.8); 3 (3.00)] | 0.63 (0.05 – 8.25) | 0.726 | 0.94 (0.07 – 12.0) | 0.963 |
| A vs. O | 25 (24.8); 52 (51.5)] | 1.33 (0.42 – 4.18) | 0.630 | 2.59 (0.84 – 8.00) | 0.099 |
| B vs. O | 21 (20.8); 52 (51.5)] | 1.68 (0.52 – 5.42) | 0.385 | 1.29 (0.34 – 4.87) | 0.703 |
| AB vs. O | 3 (3.00); 52 (51.5)] | 2.10 (0.17 – 25.5) | 0.560 | 2.75 (0.22 – 34.0) | 0.431 |
| B vs. AB | 21 (20.8); 3 (3.00)] | 0.80 (0.06 – 10.6) | 0.865 | 0.47 (0.03 – 6.57) | 0.575 |
| A vs. non-A | 25 (24.8); 76 (75.2)] | 1.10 (0.38 – 3.18) | 0.866 | 2.28 (0.81 – 6.39) | 0.117 |
| B vs. non-B | 21 (20.8); 80 (79.2)] | 1.48 (0.50 – 4.40) | 0.478 | 0.87 (0.26 – 2.94) | 0.825 |
| AB vs. non-AB | 3 (3.00); 98 (97.0)] | 1.73 (0.15 – 20.0) | 0.662 | 1.95 (0.17 – 22.6) | 0.593 |
| O vs. non-O | 52 (51.5); 49 (48.5)] | 0.66 (0.26 – 1.68) | 0.384 | 0.50 (0.19 – 1.35) | 0.172 |
| Rh blood type |  |  |  |  |  |
| Neg vs. Pos | 7 (6.90); 94 (93.1)] | 10.6 (1.89 – 58.9) | **0.007** | 6.04 (1.24 – 29.5) | **0.026** |
| ABO/Rh blood group comparison |  |  |  |  |  |
| A+ vs. A- | 22 (21.8); 3 (3.0)] | 0.11 (0.01 – 1.55) | 0.102 | 0.19 (0.01 – 2.47) | 0.203 |
| A+ vs. B+ | 22 (21.8); 20 (19.8)] | 0.67 (0.15 – 2.94) | 0.592 | 2.13 (0.45 – 9.96) | 0.339 |
| A+ vs. B- | 22 (21.8); 1 (1.00)] | 0.0 (0.0 – 0.0) | 1.000 | 0.0 (0.0 – 0.0) | 1.000 |
| A+ vs. AB+ | 22 (21.8); 3 (3.00)] | 0.44 (0.03 – 6.19) | 0.546 | 0.75 (0.06 – 9.87) | 0.827 |
| A+ vs. O+ | 22 (21.8); 49 (48.5)] | 1.14 (0.30 – 4.27) | 0.847 | 2.25 (0.66 – 7.72) | 0.197 |
| A+ vs. O- | 22 (21.8); 3 (3.00)] | 0.11 (0.01 – 1.55) | 0.102 | 0.75 (0.06 – 9.87) | 0.827 |
| A- vs. B+ | 3 (3.0); 20 (19.8)] | 6.00 (0.44 – 81.2) | 0.178 | 11.3 (0.77 – 168) | 0.078 |
| A- vs. B- | 3 (3.0); 1 (1.00)] | 0.0 (0.0 – 0.0) | 1.000 | 0.0 (0.0 – 0.0) | 1.000 |
| A- vs. AB+ | 3 (3.0); 3 (3.00)] | 4.00 (0.13 – 119) | 0.423 | 4.00 (0.13 – 119) | 0.423 |
| A- vs. O+ | 3 (3.0); 49 (48.5)] | 10.3 (0.83 – 127) | 0.070 | 12.0 (0.96 – 151) | 0.054 |
| A- vs. O- | 3 (3.0); 3 (3.00)] | 1.00 (0.03 – 29.8) | 1.000 | 4.44 (0.13 – 119) | 0.423 |
| B+ vs. B- | 20 (19.8); 1 (1.00)] | 0.0 (0.0 – 0.0) | 1.000 | 0.0 (0.0 – 0.0) | 1.000 |
| B+ vs. AB+ | 20 (19.8); 3 (3.00)] | 0.67 (0.05 – 9.02) | 0.760 | 0.35 (0.02 – 5.23) | 0.449 |
| B+ vs. O+ | 20 (19.8); 49 (48.5)] | 1.71 (0.48 – 6.05) | 0.406 | 1.06 (0.25 – 4.58) | 0.939 |
| B+ vs. O- | 20 (19.8); 3 (3.00)] | 0.17 (0.01 – 2.26) | 0.178 | 0.35 (0.02 – 5.23) | 0.449 |
| AB+ vs. B- | 3 (3.00); 1 (1.00)] | 0.0 (0.0 – 0.0) | 1.000 | 0.0 (0.0 – 0.0) | 1.000 |
| AB+ vs. O+ | 3 (3.00); 49 (48.5)] | 2.56 (0.21 – 31.8) | 0.464 | 3.00 (0.24 – 37.7) | 0.395 |
| AB+ vs. O- | 3 (3.00); 3 (3.00)] | 0.25 (0.01 – 7.45) | 0.423 | 1.00 (0.03 – 29.8) | 1.000 |
| O+ vs. B- | 49 (48.5); 1 (1.00)] | 0.0 (0.0 – 0.0) | 1.000 | 0.0 (0.0 – 0.0) | 1.000 |
| O- vs. B- | 3 (3.00); 1 (1.00)] | 0.0 (0.0 – 0.0) | 1.000 | 0.0 (0.0 – 0.0) | 1.000 |
| O+ vs. O- | 49 (48.5); 3 (3.00)] | 0.10 (0.01 – 1.21) | 0.070 | 0.33 (0.03 – 4.19) | 0.395 |

Bold numbers mean that results were statistically significant for univariate logistic analysis (p<0.05)
Abbreviations: OR, Odds ratio; CI, confidence interval

## Discussion

Generally, viral infectious diseases are a source of mortality and morbidity with significant impacts on human health around the world. SARS-CoV-2 infection remains a critical public health threat. Due to the growing number of victims related to the COVID-19 pandemic, numerous efforts have been made to identify biological factors able to influence the course of infection among COVID-19 patients. [16] It is worth mentioning that the identification of predictive biomarkers of hospitalization among patients infected with SARS-CoV-2 is essential to guide and improve clinical practice as well as reduce healthcare costs during the COVID-19 pandemic scenario. To the best of our knowledge, this is the first study that describes the impact of the blood group polymorphisms (ABO/Rh) on susceptibility and severity among COVID-19 patients in Luanda, a country located in central Africa. In this study, blood group O and Rh-positive had higher susceptibility to SARS-CoV-2 infection, severity, hospitalization, and mortality. Another study conducted by our research team also observed high susceptibility to hypertension among individuals of the O and Rh+ blood groups, showing that individuals of these blood groups might have a high susceptibility to the disease.

[17] With this study, we intend to help the clinical team in ongoing efforts to reduce hospitalization and unfavorable clinical outcomes among COVID-19 patients in Angola. Similar to our results, other studies carried out among COVID-19 patients from China, have reported several risk factors, such as older age, male gender, and the presence of underlying chronic comorbidities with SARS-CoV-2 positivity and/or unfavorable clinical outcome among COVID-19 patients. [18,19] Also, in line with prior studies carried out by our research team in Angola [20,21], we observed an increase in SARS-CoV-2 infection rate with increasing age, men were the most affected, and the urbanized areas with the highest rate of infection (Table 1).

The different categories of the ABO/Rh blood group are known to influence susceptibility to other infectious agents, such as SARS-CoV-1, where studies observed associations with the ABO/Rh blood groups. [4–6] Indeed, previous studies have shown that ABO antigen is a highly carbohydrate-enriched epitope that is present in erythrocytes, endothelial cells, and other specialized tissues that could induce a potent immune response, triggering isoagglutinin antibodies against non-expressed ABO antigens. [13,22] Spike proteins of SARS viruses are also enriched with carbohydrates, as well as ABO antigens borrowed from the SARS-infected host. [23] In this way, studies have suggested that blood group O individuals, whose blood naturally contains anti-A and anti-B isoagglutinin antibodies, have an inherent immunological advantage against SARS viral infections. [24] Even so, previous studies revealed an elevated interleukin 6 (IL-6) level in blood group O subjects compared to blood group non-O subjects. [25] This increase in IL-6 could promote the release of acute-phase proteins, such as C-reactive protein, showing that blood group O patients might experience a bad prognosis, need hospitalization, and could have an unfavorable clinical outcome [26]. Indeed, we found that O blood group patients had a higher risk of symptomatic disease (OR: 1.33, p=0.630), and hospitalization (OR: 2.59, p=0.099), compared to non-O blood group patients (Table 2), however not statistically significant. Regarding mortality, blood group O also had a high mortality rate (80%) compared to the non-O blood groups (Table 1). Niles et al. [24], also observed a higher positivity rate and worsening of SARS-CoV-2 infection among individuals with blood group O than in those non-O blood groups. However, these findings are inconsistent with those observed by Rahim et al. [5], Cheng et al. [7], Zietz et al.[9], Zhang et al. [27], and El-Shitany [26], where blood group O was less common among COVID-19 patients. Our findings also contradict the studies carried out in France (blood group A had a high risk of infection and worsening infection), Canada (blood groups A and AB had high disease severity) [28], Turkey (blood group O had lower disease severity) [29], China (blood group A had a high risk of disease severity and blood group O had low risk) [30], China (blood group A had a high risk of infection) [31], US and Denmark (no association between ABO and disease severity) [32,33], Iraq (blood group A had a high risk of disease severity) [34], and India (blood group O had low severity while blood group B had high severity). [35] The susceptibility to SARS-CoV-2 infection observed in these studies could be explained by racial, regional, and possible genetic variations.

[22] Another reason for the contradictory findings compared to our findings could be the fact that we have a homogeneous sample since the Angolan population is mostly from the blood group O and Rh-positive, which could suggest false protection to SARS-CoV-2 among the non-O blood group. Indeed, previous studies documented that in Angola, blood group O represents 54.4% of the population, followed by blood groups A (22.3%), B (19.7%), and blood group AB is the least frequent with 3.7%. [36] Therefore, comparing the general frequency of blood groups in the healthy Angolan population with the SARS-CoV-2 positive population, there was a reduction in the frequency of blood groups O (54.4% to 51.5%) and AB (3.7% to 3.0%), while an increase in the frequency was observed in blood groups A (22.3% to 24.8%) and B (19.7% to 20.8%) (Tables 1 and 2). These results could indicate that non-O or blood groups A and B are the ones with the highest risk for SARS-CoV-2 infection in Angola. Indeed, this highest risk is consistent with our findings, since 24.4% and 19.2% of COVID-19 patients in groups A and B, respectively, showed symptoms related to the SARS-CoV-2 infection. In addition, 21.3% of patients in groups A and B, simultaneously, were hospitalized due to the worsening of their clinical condition, and 20% of patients in blood group A died due to COVID-19 (Table 1). Similar to the study carried out by Zietz et al. [9], only blood group B had inconsistent effects between the risk of developing symptomatic SARS-CoV-2 infection and hospitalization. Both studies observed that patients in blood group B, despite having a lower risk of developing symptomatic infection (OR: 0.79, p=0.725), are more likely to be hospitalized (OR: 2.0, p=0.323) due to worsening of SARS-CoV-2 infection (Table 2). At this time, we do not have a reasonable explanation related to the need for hospitalization among blood group B patients, however, further studies need to be conducted. Even so, it is worth mentioning that a meta-analysis, carried out by Dentali et al. [37], found that the non-O blood group is a candidate to be one of the most important genetic risk factors for venous thrombosis. Although non-O patients (48.5%, 49/101) were the least frequent in our studied population, coagulopathy [38,39] and/or the risk of venous thromboembolism [40,41] must be evaluated since these hematological disorders have been reported to be a common issue for COVID-19 patients.

Differences in the risk of SARS-CoV-2 infection were also observed among the Rh blood types. The Rh- positive patients presented a high rate of symptomatic infection compared to Rh-negative patients (78.3% to 21.7%, p=0.001) (Table 1). These differences were also observed by Niles et al. [24], who showed that Rh positivity, regardless of the ABO blood group, was a significant risk factor for SARS-CoV-2 infection. Furthermore, the risk of developing symptomatic infection (OR: 10.6, p=0.007) and need for hospitalization (OR: 6.04, p=0.026) was high among patients with Rh-positive blood type compared to Rh-negative patients (Table 2). These results are in contrast to that reported in Pakistan where the likelihood of Rh-positive blood types to be SARS-CoV-2 positive was 0.75 (95% CI 0.57-0.98) [5] but are similar to that observed among COVID-19 patients from New York [9] and in the state of Massachusetts. [13] On the other hand, Abdollahi et al. [6], have observed no relationship between Rh blood type and susceptibility to SARS-CoV-2 infection, showing that the association between Rh-positive blood type and risk of developing symptomatic SARS-CoV-2 infection or need for hospitalization observed in the present study, merits further investigation.

In the present study, the most common symptoms among COVID-19 patients were cough (36.6%), fever (35.6%), asthenia (26.7%), malaise (19.8%), dyspnoea (18.8%), and headache (14.9%) (Table 1). These symptoms were in accordance with those reported in previous studies. [3,26] However, our findings emphasize the need for higher attention to the relationship between fever, malaise, and anorexia, with ABO blood groups (p<0.05), mainly in COVID-19 patients from the AB blood group, since 10% and 16.7% of these patients had malaise and anorexia, respectively (Table 1). The reasons for this relationship more frequently in the AB group are not understood and need further investigation. It is also worth mentioning that despite being uncommon, the patients in the blood group O were the only ones who presented hemiplegia and loss of consciousness, which needs to be explored in future studies. On the other hand, all COVID-19 patients who died were of Rh-positive blood type (Table 1). Interestingly, the AB blood group was the only blood group that showed a significant relationship with malaria (33.3%, p=0.002), a vector-borne disease (VBD) endemic in Luanda, the capital city of Angola (Table 1). Therefore, studies on the relationship between SARS-CoV-2 and VBD such as malaria, dengue, zika, chikungunya, and yellow fever, should be urgently carried out, as there have been outbreaks of VBD in Angola. [42–44] Indeed, a study recently carried out by our research team observed a coinfection rate between SARS-CoV-2 and VBD of 11.4%, of which, 14.3% of patients were coinfected with malaria and 10.3% were coinfected with dengue, suggesting that patients with COVID-19 should also be screened for VBD and vice versa. [45]

Our findings might have a positive implication for clinicians and policymakers, especially in the categorization of patients regarding susceptibility to SARS-CoV-2 infection, severity, hospitalization, and mortality according to the ABO blood group of the individuals. At an early stage, we can present a description of the Angolan individuals as follows: First, the individuals in blood group A tend to present moderate susceptibility, severity, hospitalization, and mortality, respectively. Second, the individuals in blood group B tend to present moderate susceptibility, low severity, high hospitalization, and low mortality. Third, the individuals in group AB tend to present a low susceptibility, severity, hospitalization, and mortality, respectively. Finally, the individuals in group O tend to present a high susceptibility, severity, hospitalization, and mortality, respectively.

This study had some potential limitations. Although being a multicentric study, the sample size might not represent whole COVID-19 patients in Luanda. Moreover, negative COVID-19 patients were not included as a control group. Despite these weaknesses, these are preliminary results from COVID-19 patients in an African country, and agreement with other published studies showed that the relationship between ABO/Rh blood groups and SARS-CoV-2 is not yet consistent, and the scientific community has yet to come up with a reasonable explanation for the relationship between ABO/Rh blood groups and SARS-CoV-2 infection. Further studies should be carried out in order to have a clearer insight into the relationship between ABO/Rh blood groups and SARS-CoV-2 severity as well as hematological, biochemical, and immunological laboratory abnormalities according to ABO/Rh blood groups among COVID-19 patients in Angola.

## Conclusion

Our findings showed a putative relationship between the ABO/Rh blood group with SARS-CoV-2 severity and hospitalization. COVID-19 patients from blood group O and Rh-positive showed a high likelihood related to SARS-CoV-2 susceptibility, severity, hospitalization, and mortality, respectively, while blood group AB presented a low susceptibility, severity, hospitalization, and mortality, respectively. Moreover, the results of this study add to the growing body of evidence suggesting that ABO/Rh blood groups play an important role in the course of SARS-CoV-2 infection.

## Acknowledgments

The authors appreciate the participation of all Angolan COVID-19 patients enrolled in the study. We would also like to express our gratitude to the Fundação Calouste Gulbenkian (FCG) and Camões, IP, for financial support. Reagents for the determination of ABO/Rh blood groups among COVID-19 patients were generously provided by the Instituto Nacional de Sangue of the Ministry of Health of Angola. Appreciation also goes to CISA, INIS, Hospital Militar Principal, Clínica Girassol, and Lucrécia Paim Maternity, for institutional support. We would also like to acknowledge Anabela Mateus, Domingos Alfredo, and Ngueza Loureiro, for laboratory support; Zinga David, António Mateus, Eunice Manico, Paolo Parimbelli, Deodete Machado, and Manuela Mendes for administrative support; Vera Mendes, Sandra Ferreira, and Joana Sebastião for logistical support.

## Data availability statement

All relevant data are within the manuscript.

## Author contributions

Conceptualization and methodology: CSS, JNV, and MB. Formal analysis and data curation: CSS and MB. Investigation: CSS, AT, AL, MA, CT, AC, and BC. Supervision: CSS, JNV, and MB. Project administration: CSS, ES, JM, JNV, and MB. Writing—original draft preparation: CSS. Writing—review and editing: CSS, JNV, JM, MA, CT, AC, ES, and MB. All authors read and approved the final manuscript.

